# Specificity Guides Interpretation: On H3K4 Methylation at Enhancers and Broad Promoters

**DOI:** 10.1101/2023.01.16.524067

**Authors:** Rohan N. Shah, Alexander J. Ruthenburg

**Author notes:** Correspondence to (R.N.S.); (A.J.R.).

## Abstract

In 2018, we used internally calibrated chromatin immunoprecipitation (ICeChIP) to find that many of the most commonly used antibodies against H3K4 methylforms had significant off-target binding, which compromised the findings of at least eight literature paradigms that used these antibodies for ChIP-seq (Shah et al., 2018). In many cases, we were able to recapitulate the prior findings in K562 cells with the original, low-quality antibody, only to find that the models did not hold up to scrutiny with highly specific reagents and quantitative calibration.

In a recent preprint originally prepared as a Letter to the Editor of Molecular Cell, though they agree with our overarching conclusions, Pekowska and colleagues take issue with analyses presented for two relatively minor points of the paper (Pekowska et al., 2023). We are puzzled by the assertion that these two points constitute the “bulk” of our findings, nor is it clear which components of our “analytical design” they find problematic. We feel their critique, however mild, is misguided.

## Main Text

Ultimately, the points that Pekowska and colleagues raise fall into two categories. First, they argue that our conclusions do not necessarily hold outside of K562 cells, a point that we ourselves repeatedly cautioned in the paper. Second, they argue that H3K4me3 is a biologically relevant marker of enhancers, which ultimately reduces to the definition of enhancers used. We contend that their definition encompasses many unannotated promoters. Even with this liberal definition, notional H3K4me3-containing enhancers represent a tiny fraction of their candidates.

Based on uncalibrated ChIP with a poorly specific antibody in CD4+ T cells, Pekowska and colleagues previously argued that H3K4me2 was enriched over immune system process gene bodies (Pekowska et al., 2010). Using the same low-quality antibody in K562 cells (Fig. 1A), we partially recapitulated this enrichment of H3K4me2 over gene bodies, but noted that the breadth of this enriched region was diminished with high-quality H3K4me2 antibodies (Shah et al., 2018). Further, we found that in K562s, genuine H3K4me2 gene body enrichment largely marked metabolic rather than immune system process genes (Shah et al., 2018). We could not determine whether these differences were driven by antibody quality, ChIP procedure, or cell type (pg. 173, Shah et al., 2018) – though the observed differences between low- and high-quality antibodies within the single cell type of K562s suggests that antibody specificity was at least a contributing factor.

**Figure 1:**
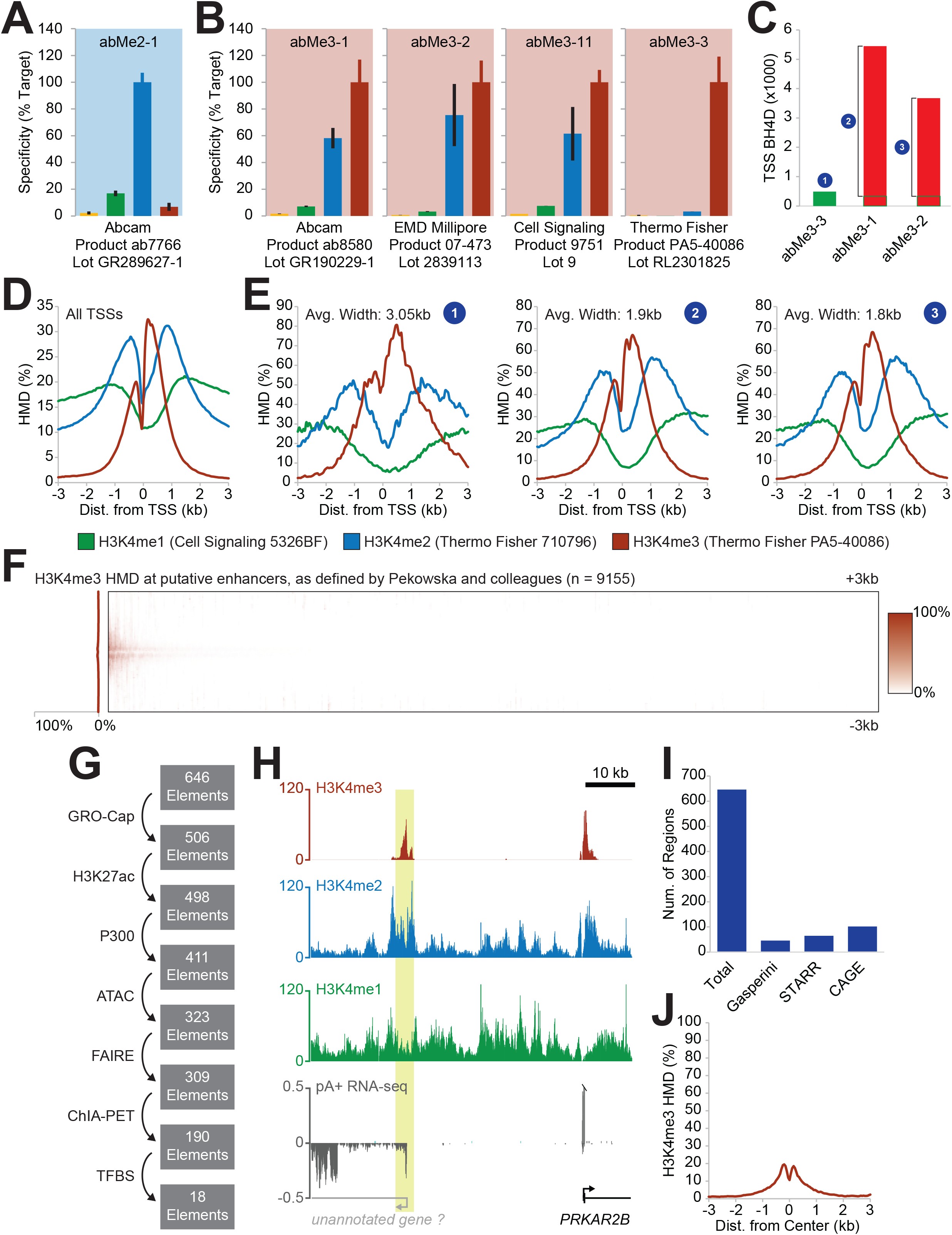
H3K4me3 Antibody Specificity and Distribution. **(A)** Specificity of H3K4me2 antibody abMe2-1, used by Pekowska et al., 2010. Adapted from Shah et al., 2018. **(B)** Specificity of low-quality H3K4me3 antibodies (abMe3-1, abMe3-2, and abMe3-11) and high-quality H3K4me3 antibody (abMe3-3). Note that, given the greater abundance of H3K4me2, the low-quality antibodies will globally capture more H3K4me2 than H3K4me3. Adapted from Shah et al., 2018. **(C)** Number of transcription start site (TSS) broad H3K4me3 domains (BH4D), defined as a contiguous region above 25% HMD intersecting the TSS with a length of at least 2kb, using a high-quality H3K4me3 antibody (abMe3-3, green) or low-quality H3K4me3 antibodies (abMe3-1 and abMe3-2, red), which also bind H3K4me2 (see panel B). Numbered blue circles correspond to subsets of TSSs presented in panel E. Green outlines represent the number of BH4D from low-quality antibodies that were found by high-quality antibody. **(D)** Metagene profile of H3K4me1, H3K4me2, and H3K4me3 histone modification density (HMD) at all TSSs using highly specific antibodies (Shah et al., 2018). **(E)** High-specificity metagene profiles of H3K4me1, H3K4me2, and H3K4me3 HMD contoured over either the set of TSSs with BH4D in highly specific abMe3-3 dataset (left), or the set of extra TSSs with BH4D in either low-specificity abMe3-1 (center) or low-specificity abMe3-2 (right) datasets that were not found in the high-specificity abMe3-3 dataset. Note similarity of (2) and (3) to panel D. **(F)** Heatmap of H3K4me3 HMD with abMe3-3 across all putative enhancers defined by Pekowska and colleagues. **(G)** Number of regions remaining after filtering putative distal enhancers with H3K4me3 peaks (per Pekowska and colleagues) through filtering steps previously employed in Shah et al., 2018. Enhancers were defined as “those not overlapping with a Refseq promoter and have a transcription factor binding site [TFBS] (Wang et al., 2014), GRO-Cap TSS (Core et al., 2014), ATAC-seq peak (Buenrostro et al., 2015), and ENCODE H3K27ac peak, FAIRE-seq peak, DNase HS site, and P300 peak (The ENCODE Project Consortium, 2012) which make contact with at least one promoter by pol II ChIA-PET (Li et al., 2012)” (Shah et al., 2018). **(H)** Genome browser view of high-specificity H3K4me1, H3K4me2, and H3K4me3 HMD with polyA+ RNA-seq from ENCODE (Davis et al., 2018; The ENCODE Project Consortium, 2012) at the putative enhancer presented by Pekowska and colleagues highlighted in yellow (their Fig. 1D; Pekowska et al., 2023). **(I)** Number of putative distal enhancers with H3K4me3 peaks (per Pekowska and colleagues) that overlap with validated K562 enhancers from Gasperini et al., 2019 (n = 46); K562 STARR-seq peaks from ENCODE (Davis et al., 2018; The ENCODE Project Consortium, 2012) (n = 65); or K562 bidirectional CAGE peaks from FANTOM5 (Hon et al., 2017) (n = 102). **(J)** Metagene profile of H3K4me3 HMD contoured over putative distal enhancers with H3K4me3 peaks (per Pekowska and colleagues; n = 646). Note the apical average histone modification density is approximately 20%.

We are pleased that Pekowska and colleagues recapitulate our findings in K562s (their Fig. 1A-B). However, none of their new analyses shed light on which factor contributed most to this discrepancy because they have not re-evaluated their findings in any other cell type with a sufficiently specific antibody. They attempt to argue that antibody quality and calibration are not relevant for this purpose because uncalibrated ChIP data from low- and high-quality antibodies yield superficially similar gene ontology results in K562s. This logic is specious; the fact that low-quality ChIP data does not fail in *all* cases does not imply that it is generally useful, particularly outside of a highly coarse-grained and aggregative analysis.

Similarly, for the H3K4me3 breadth analyses, though we agree that cell types may account for some differences with the literature, we note that the role of broad H3K4me3 domains (BH4D) was previously published in K562 cells using a low-quality antibody (Chen et al., 2015) – a paradigm that we showed to be inaccurate in that same cell line with high-specificity reagents (Shah et al., 2018). No extant analyses definitively deconvolute the contributions of antibody quality and cell type to these findings, so it is difficult to interpret the other H3K4me3 datasets (their Fig. 1C) without calibration, particularly given that those antibodies have 50-70% cross-reactivity with H3K4me2 (Fig. 1B), a far more abundant modification (LeRoy et al., 2013). These antibodies are particularly poor for analyses of BH4Ds; because H3K4me2 flanks a central peak of H3K4me3 around promoters, off-target capture of the former expands the breadth of the apparent enriched region of the latter, resulting in many false-positive BH4Ds (Fig. 1C-E). And even if there are genuine differences in H3K4me3 breadth between cell lines, we do not see those results as conflicting with ours; as we noted repeatedly in our paper (Shah et al., 2018), our findings were limited to K562s and only directly refutes models in that cell line (Chen et al., 2015).

Pekowska and colleagues also address the question of enhancer H3K4me3. Their previous work used low-quality antibodies to argue that H3K4me3 marks active enhancers (Pekowska et al., 2011). In their preprint, they cite six additional papers in support of this idea; however, these papers rely exclusively on one or more of the three low-specificity antibodies presented in Fig. 1B, which are incapable of distinguishing H3K4me3 from H3K4me2. We recapitulated this apparent H3K4me3 signal with low-quality antibodies in K562s at a stringently defined set of enhancers (Fig. 5B-E, Shah et al., 2018). We then showed that there is actually very little H3K4me3 at these enhancers, with apparent H3K4me3 signal from low-quality antibodies being driven by off-target capture of H3K4me2. In their preprint, to support their prior conclusions, Pekowska and colleagues point to Figure 4H in our paper (Shah et al., 2018), claiming that it shows that there is a “significant level of H3K4me3 at enhancers” (Pekowska et al., 2023). They also state that 6% of distal enhancers in K562s have H3K4me3 peaks, which they claim constitutes a genuine mark of enhancer activity based on correlation with other findings, some of which we conclusively refuted (Koch et al., 2011; Shah et al., 2018).

We feel their analysis is problematic in several respects. First, the figure panel to which they point as showing “a significant level of H3K4me3 at enhancers” shows that on average, only 1-5% of nucleosomes would bear H3K4me3, a quantity sufficiently little to be of questionable biological significance. By contrast, at highly transcribed promoters, this quantity approaches 100% (Grzybowski et al., 2015; Shah et al., 2018).

More worrisome, Pekowska and colleagues employ an enhancer definition that is exceedingly broad and does not effectively exclude poorly annotated promoters. In our study, we used very stringent criteria to define enhancers to avoid conflating the methylation states at promoters with those at enhancers, particularly given the strong association between promoters and H3K4me3 (Fig. 1G). By contrast, Pekowska and colleagues define enhancers as regions with DNase HS sites and an H3K4me1 peak not overlapping an annotated Gencode promoter, which is considerably more permissive to unannotated promoters. Their genome browser views (their Fig. 1D) demonstrate this permissiveness; the top example does not meet their criteria because the H3K4me3 peak and DHS don’t overlap, and the bottom appears to be a unidirectional promoter for a polyadenylated long RNA (Fig. 1H). Even with this exceedingly lax definition of “enhancers,” heatmap analysis of their enhancer set reveals H3K4me3 to be minimal at these regions (Fig. 1F).

Attempting to reproduce their analysis with their code, we find 646 possible “enhancers” that overlap an H3K4me3 peak (curiously, 109 more than they identify with the same code). However, going through each of the filtering steps listed above, we find that only 18 such loci would meet our criteria for stringently defined enhancers (Fig. 1G), representing approximately 0.1% of their loosely defined enhancer set (18/9155). If this set is further compared against the FANTOM5 list of validated enhancers (Andersson et al., 2014) and a set of unannotated lncRNA promoters (Iyer et al., 2015), there is only *one* locus that would be classified as a distal enhancer with an H3K4me3 peak. Other functional enhancer definitions have similarly low overlap with this subset of candidates (Fig. 1I). We question whether this tiny subset is biologically relevant; we would argue that these rare loci are, at best, exceptions that should not be intermingled with more pervasive enhancer features.

Further, we would argue that peak calling is inadequate for identifying regions with functionally relevant amounts of histone modifications. The fact that there is more signal than background (i.e., a called peak) does not inherently imply that there is a biologically relevant quantity of the histone modification at that region. Given the high sequencing depth we employ with our ICeChIP datasets, we routinely find called peaks wherein <1% of nucleosomes at that region have the histone modification of interest (Grzybowski et al., 2015; Shah et al., 2018). This can be seen even in the loci selected by Pekowska and colleagues; in their Fig. 1D, there is an example of a region that has a high-confidence H3K4me3 peak but very little H3K4me3 enrichment. Overall, at the regions identified by Pekowska and colleagues as being H3K4me3-marked, only a small minority of these loci appear to be distinguishable from promoters, and of those loci, only a small minority of the nucleosomes actually bear H3K4me3 (Fig. 1J), making its functionality questionable.

Given both the very loose definition of enhancers and the low level of H3K4me3 present at even this very permissive subset, it is difficult to sustain an assertion of H3K4me3 being a significant and genuine mark of enhancers in K562s. Pekowska and colleagues try to justify this assertion with correlation to other datasets (their Fig. 1E-F), but they use features that are common between promoters and enhancers (GRO-Cap TSSs, Pol II peaks, etc.), or features that have been shown to not be associated with enhancers (e.g., intergenic TBP sites; Shah et al., 2018), rendering those conclusions irrelevant.

We greatly appreciate that Pekowska and colleagues agree with the need for the higher standards of ChIP-seq that ICeChIP can provide. However, their arguments here still make the error of trying to infer biological meaning from ChIP experiments with high off-target capture.

## Acknowledgements

This study was supported by the National Institutes of Health under award numbers R01-GM115945-05 and R01-HL148719 to A.J.R. and T32-HD007009-45 for training grant funding provided to R.N.S. The funders had no role in study design, data collection and analysis, decision to publish, or preparation of the manuscript.

## Software and Code Availability

R code for these analyses can be found at https://github.com/shah-rohan/mw-pekowska2019.

## Methods and Materials

### ICeChIP-seq Datasets

ICeChIP-seq datasets were previously published (Shah et al., 2018) and downloaded from the Gene Expression Omnibus, listed under accession number GSE103543.

### Broad H3K4me3 Domain Analysis

Broad H3K4me3 domains were defined as a contiguous region with a length of at least 2000bp wherein the apparent H3K4me3 HMD was greater than or equal to 25% in consecutive 50bp windows and such that this domain intersects the annotated transcription start site. This was accomplished using the breadth.R script provided in the GitHub repository under “Software and Code Availability.”

### CAGE Dataset Analysis

CAGE peaks mapped to hg38 were obtained from FANTOM5 (Hon et al., 2017) from the dataset with the library ID CNhs11250. The set of CAGE peaks was split in to positive and negative strand files, extended by 400bp, and intersected with BEDTools (Quinlan and Hall, 2010) to identify sites with opposite strand CAGE peaks within 400bp of each other. The region between corresponding peaks was then treated as the region of bidirectional CAGE TSSs for the purposes of intersection with the set of putative enhancers defined by Pekowska and colleagues.

### STARR-seq Dataset Analysis

Two replicates of STARR-seq peaks in K562 were obtained from ENCODE (Davis et al., 2018; The ENCODE Project Consortium, 2012), accession numbers ENCFF394DBM and ENCFF717VJK. BEDTools was used to identify peaks that were present in both these replicates; these were treated as the set of STARR-seq peaks for intersection with the set of putative enhancers defined by Pekowska and colleagues.

### Gasperini Enhancer Dataset Analysis

Validated K562 enhancers were obtained from Gasperini et al., 2019, then lifted over from hg19 to hg38 for intersection with the set of putative enhancers defined by Pekowska and colleagues.

### RNA-seq Dataset Analysis

RNA-seq datasets were obtained from ENCODE (Davis et al., 2018; The ENCODE Project Consortium, 2012), accession numbers ENCFF887YZQ and ENCFF203XQW.

### Metagene Profiles

Metagene profiles were generated using HOMER with the *annotatePeaks*.*pl* tool as previously described (Shah et al., 2018).

### Filtering Analysis

Putative enhancers defined by Pekowska and colleagues were filtered for the presence of a transcription factor binding site (Wang et al., 2014), GRO-Cap TSS (Core et al., 2014), ATAC-seq peak (Buenrostro et al., 2015), and ENCODE H3K27ac peak, FAIRE-seq peak, and P300 peak (Davis et al., 2018; The ENCODE Project Consortium, 2012), with contact to at least one promoter by pol II ChIA-PET (Li et al., 2012), as previously done (Shah et al., 2018). Further filtering was done with FANTOM5 K562 enhancers (Andersson et al., 2014) and an annotation of lncRNAs in K562s (Iyer et al., 2015) in a modified version of the R code provided by Pekowska and colleagues. This modified version is provided in the GitHub repository under “Software and Code Availability.”

### Summary of Datasets Used for This Study

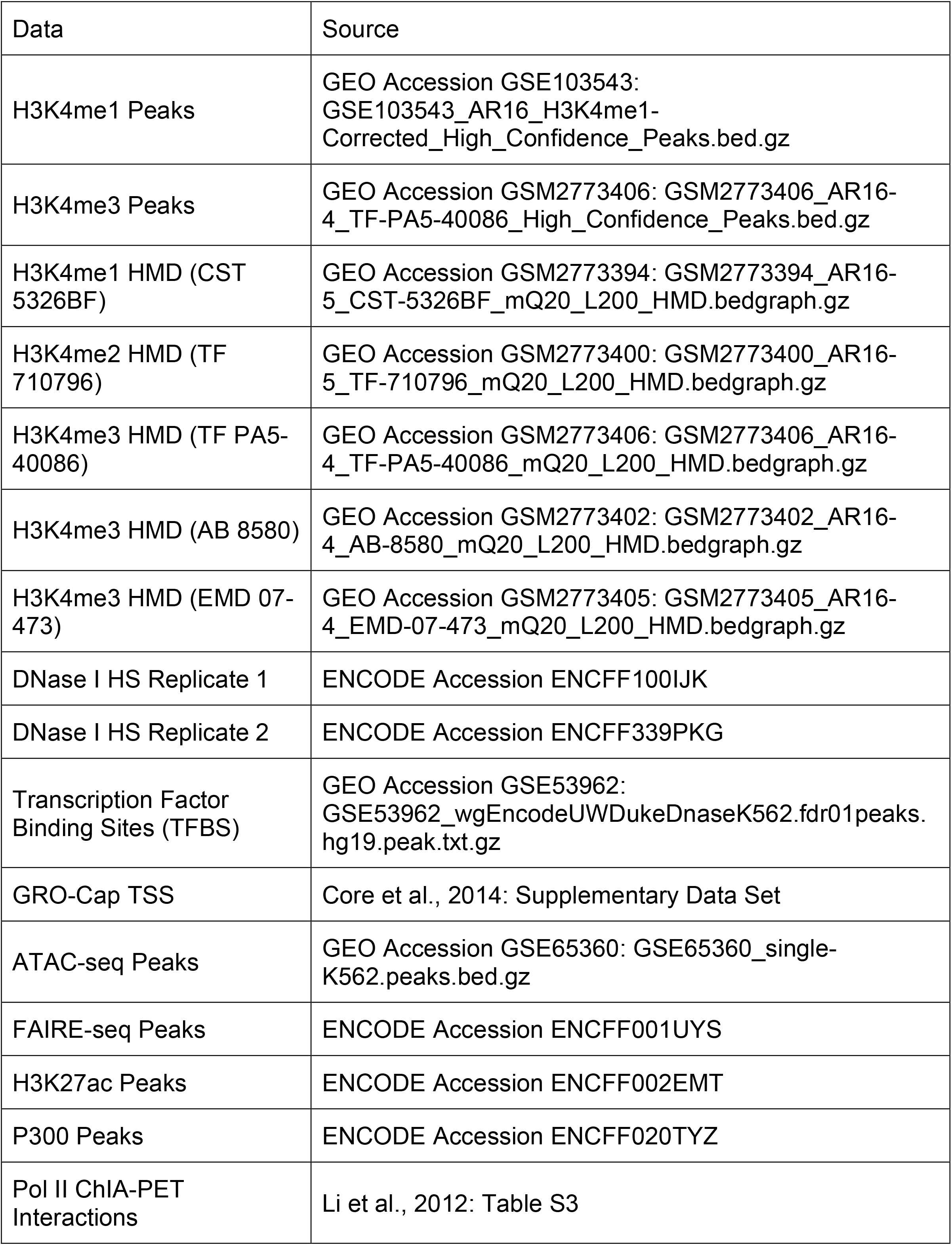
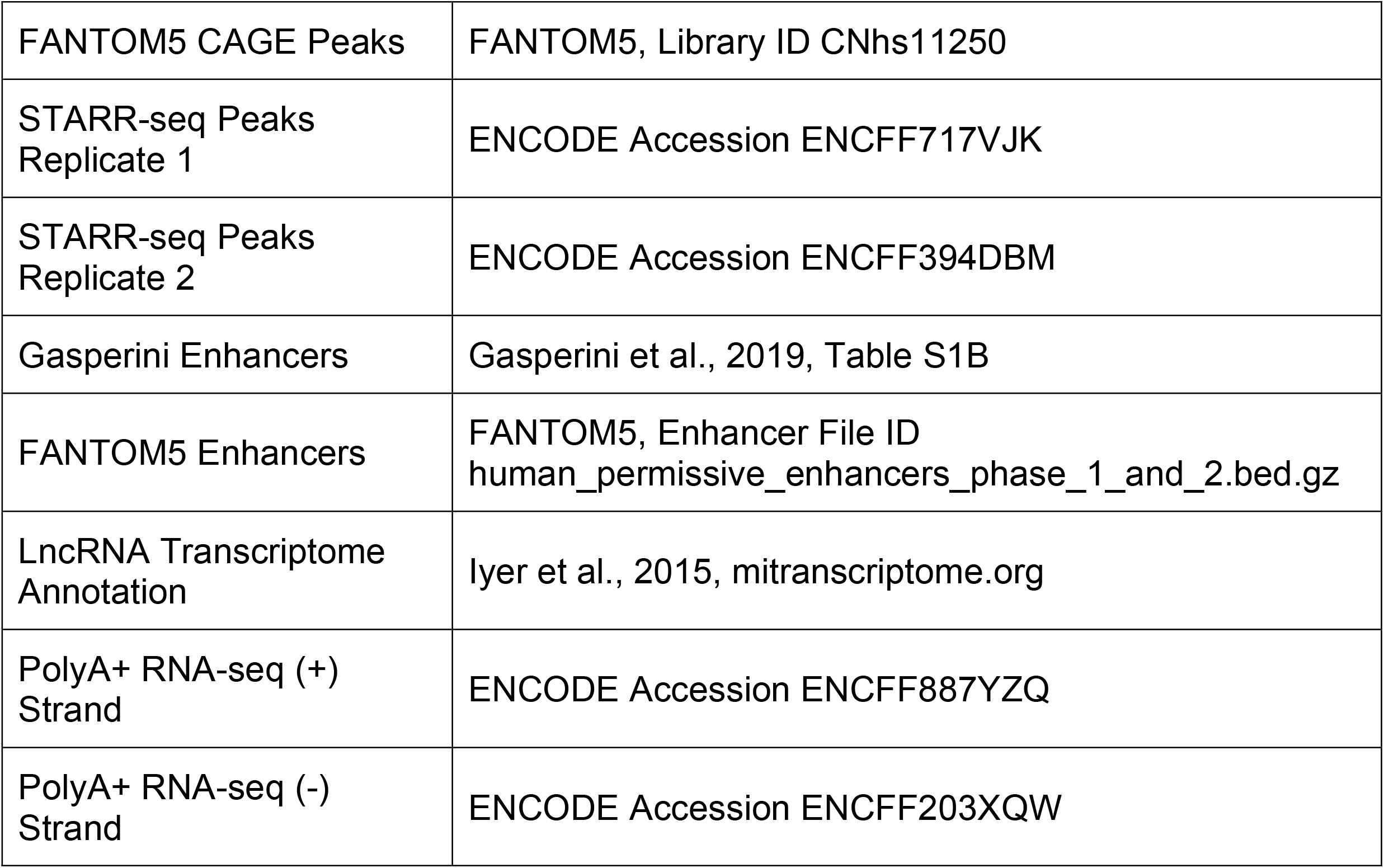

